# *Spodoptera littoralis* genome mining brings insights on the dynamic of expansion of gustatory receptors in polyphagous noctuidae

**DOI:** 10.1101/2021.12.09.471925

**Authors:** Camille Meslin, Pauline Mainet, Nicolas Montagné, Stéphanie Robin, Fabrice Legeai, Anthony Bretaudeau, J. Spencer Johnston, Fotini Koutroumpa, Emma Persyn, Christelle Monsempès, Marie-Christine François, Emmanuelle Jacquin-Joly

## Abstract

The bitter taste, triggered via gustatory receptors, serves as an important natural defense against the ingestion of poisonous foods in animals, and the diversity of food diet is usually linked to an increase in the number of gustatory receptor genes. This has been especially observed in polyphagous insect species, such as noctuid species from the *Spodoptera* genus. However, the dynamic and physical mechanisms leading to these gene expansions and the evolutionary pressures behind them remain elusive. Among major drivers of genome dynamics are the transposable elements but, surprisingly, their potential role in insect gustatory receptors expansion has not been considered yet.

In this work, we hypothesized that transposable elements and possibly positive selection would be involved in the active dynamic of gustatory receptor evolution in *Spodoptera spp*. We first sequenced *de novo* the full 465Mb genome of *S. littoralis*, and manually annotated all chemosensory genes, including a large repertoire of 373 gustatory receptor genes (including 19 pseudogenes). We also improved the completeness of *S. frugiperda* and *S. litura* gustatory receptor repertoires. Then, we annotated transposable elements and revealed that a particular category of class I retrotransposons, the SINE transposons, was significantly enriched in the vicinity of gustatory receptor gene clusters, suggesting a transposon-mediated mechanism for the formation of these clusters. Selection pressure analyses indicated that positive selection within the gustatory receptor gene family is cryptic, only 7 receptors being identified as positively selected.

Altogether, our data provide a new good quality *Spodoptera* genome, pinpoint interesting gustatory receptor candidates for further functional studies and bring valuable genomic information on the mechanisms of gustatory receptor expansions in polyphagous insect species.

## Introduction

Animals rely heavily on their sense of taste to discriminate between harmful poisonous foods, usually through the detection of bitter taste, and beneficial sustenance. Interestingly, narrowness of food diets in animals is usually linked to a decreased number of gustatory receptors (GRs), in both mammals such as the blood-feeder bats^1^, and in insects such as the body louse^2^ - an obligate ectoparasite of human -, the fig wasp *Ceratosolen solmsi*^3^– specialized on *Ficus* – and many Lepidoptera specialist feeders, although mammals and insect GRs are unrelated. Reversely, the diversity of food diet is usually linked to GR gene expansions. This has been especially observed in polyphagous insects, including omnivorous species such as the American cockroach *Periplaneta americana*^4^ and herbivorous species such as noctuid species^5–7^.

In polyphagous noctuids, the sequencing of the genomes of *Spodoptera frugiperda* and *S. litura* revealed GR repertoires of 231 and 237 genes^5, 8^, respectively, more than twice as much compared to other monophagous and oligophagous Lepidoptera species (*Bombyx mori*: 69 genes, *Heliconius melpomene*: 73 genes)^9–12^, suggesting that the number of GRs has greatly increased during evolution in polyphagous Lepidoptera via gene tandem duplication. The genomic architecture of the GR family is thus well known in these species and, together with previous studies, it supports the evidence that the family evolved under a birth-and-death model as well as under different selective pressures depending on the clade considered^10, 13–15^. Most of these GRs belong to clades grouping the so-called “bitter” receptors, but in fact the function of the majority of these GRs remains enigmatic. Although the bitter GR class exhibits the most dynamic evolution, the mechanisms leading to GR expansions and the evolutionary pressures behind them remain elusive. Among major drivers of genome dynamics are the transposable elements (TEs). TEs are very diverse and are distributed along genomes in a non-random way^16^. Similar or identical TEs can induce chromosomal rearrangements such as deletions, insertions and even duplications ^17–19^, features that are frequent in multigene families such as GRs. Surprisingly, their potential role in insect GR expansion has not been considered yet. In order to study in more details GRs evolution and the potential role of TEs in GRs expansion, we sequenced an additional genome of a Spodoptera species: *Spodoptera littoralis.* So far, only 38 GRs identified^20–22^ in *S. littoralis* whereas several hundreds of GRs were annotated in its counterparts *S. litura* and *S. frugiperda*. To investigate this singularity, we report here the sequencing of the *S. littoralis* genome, its full assembly, functional automatic annotation and expert annotation of all chemosensory gene families, namely soluble carrier proteins (odorant-binding proteins: OBPs, and chemosensory proteins: CSPs)^23^ and the three major families of insect chemosensory receptors (odorant receptors: ORs, ionotropic receptors: IRs and GRs)^24^. With a particular focus on gustation, we also reannotated GRs in *S. litura* and *S. frugiperda*. Then, we analyzed the evolutionary history of GRs, by looking at the enrichment for transposable elements in the vicinity of GRs and by analyzing selective pressures acting on the different GR clades.

## Methods and Materials

### Estimation of *Spodoptera littoralis* genome size

The genome size of *Spodoptera littoralis* was estimated using flow cytometry. Genome size estimates were produced as described before^25^. In brief, the head of a *S. littoralis* along with the head of a female *Drosophila virilis* standard (1C = 328 Mbp) were placed into 1 ml of Galbrath buffer in a 2 ml Kontes Dounce and ground with 15 strokes of the A pestle. The released nuclei were filtered through a 40 μM nylon filter and stained with 25 μg/mL propidium iodide for 2 hours in the cold and dark. The average red fluorescence of the 2C nuclei was scored with a Partec C flow cytometer emitting at 514 nm. The 1C genome size of *S. littoralis* was estimated as (average red florescence of the 2C *S. littoralis* peak) / (average fluorescence of the 2C *D. virilis* peak) X 328 Mbp.

### *Spodoptera littoralis* genome sequencing and assembly

#### Biological material and genomic DNA extraction

Whole genomic DNA was extracted from two male larvae obtained after two inbred generations resulting from a single pair of *S. littoralis* originating from a laboratory colony maintained in INRAE Versailles since 2000s on suitable laboratory diet (Poitout and Bues 1974). The sex of individuals was verified by checking for presence of testis. The gut was removed and DNA extraction was performed from whole, late-stage larvae using Qiagen Genomic-tip 500/G (Qiagen Inc., Chatsworth, CA, USA). A total of 30 µg of genomic DNA was obtained.

#### Sequencing

Different types of libraries were generated for two sequencing technologies: Illumina and PacBio. For Illumina sequencing, five libraries were prepared and constructed according to the Illumina manufacturer’s protocol (one library of 170, one of 250 and three of 500 bp). Illumina sequencing was performed at the BGI-tech facilities (Shenzen, China) on a HiSeq2500 machine. Around 68 Gb were obtained, representing 144X of the estimated genome size (470 Mb) (Supp data 1). The raw reads were filtered at BGI to remove adapter sequences, contaminations and low-quality reads and the quality of all raw reads was assessed using FASTQC (Andrews S. http://www.bioinformatics.babraham.ac.uk/projects/fastqc/). PacBio sequencing was performed at GenoScreen (Lille, France) by the SMRT sequencing technology on 9 SMRTcell RSII, generating 2 846 820 reads. Around 16 Gb were obtained, representing 34X of the estimated genome size (Supp data 1). High quality sequences were obtained by generating circular consensus sequencing (CCS).

#### Genome assembly

A first assembly was done using Platanus (v1.2.1)^26^ with Illumina data. A second assembly was obtained by doing scaffolding with SSPACE-LR (modified)^27^ using PacBio data and gap filling using GapCloser^28^. These second assembly was evaluated using Benchmarking Universal Single-Copy Orthologue (BUSCO v3.0.2)^29^ with a reference set of 1,658 genes conserved in Insecta.

### Structural and functional genome annotation

Structural automatic genome annotation was done with BRAKER (v1.11) ^30^ using all RNAseq data described in Supp data 1. RNAseq libraries were sequenced from different larvae and adult tissues from males and females including the proboscis, palps, legs and ovipositor and sequenced by Illumina (Supp data 1)^20–22^. Reads were trimmed using Trimmomatic (v0.36)^31^ with the following parameters : ILLUMINACLIP:TruSeq2-PE.fa:2:30:10, LEADING:3, TRAILING:3, SLIDINGWINDOW:4:15, MINLEN:36. Trimmed reads were mapped on the genome assembly using STAR (v.5.2a)^32^ with the default parameters except for the following parameters : outFilterMultimapNmax = 5, outFilterMismatchNmax = 3, alignIntronMin = 10, alignIntronMax = 50000 and alignMatesGapMax = 50000. As done for the genome assembly, gene annotation was evaluated using Benchmarking Universal Single-Copy Orthologue (BUSCO v3.0.2) ^29^ with a reference set of 1,658 proteins conserved in Insecta. Putative functions of predicted proteins were assigned using blastp (v2.6.0) against GenBank NR (non-redundant GenBank CDS translations+PDB+SwissProt+PIR+PRF) release 09/2017, and interproscan v5.13-52.0 against Interpro. Associated GO terms were collected from blast NR and interproscan results with blast2GO (v2.5).

### Annotation of OBPs, CSPs, ORs and IRs

The annotation of genes encoding soluble transporters (OBPs and CSPs), odorant receptors (ORs) and ionotropic receptors (IRs) was performed using known sequences from other species with their genome sequenced (*S. frugiperda*, *S. litura*, *B. mori*, *H. melpomene* and *Danaus plexippus*)^8, 10, 11, 33, 34^. For each type of gene family, the set of known amino acid sequences and the genome sequence of *S. littoralis* were uploaded on the BIPAA galaxy platform to run the following annotation workflow. First, known amino acid sequences were used to search for *S. littoralis* scaffolds potentially containing genes of interest using tblastn^35^. All *S. littoralis* scaffolds with significant blast hits (e-value < 0.001) were retrieved to generate a subset of the genome. Amino acid sequences were then aligned to this subset of the genome using Scipio^36^ and Exonerate^37^ to define intron/exon boundaries and to create gene models. Outputs from Scipio and Exonerate were then visualized on a Apollo browser^38^ available on the BIPAA platform. All gene models generated have been manually validated or corrected via Apollo, based on homology with other lepidopteran sequences and on RNAseq data available for *S. littoralis*^20, 22, 39^. The classification of deduced proteins and their integrity were verified using blastp against the non-redundant (NR) GenBank database. When genes were suspected to be split on different scaffolds, protein sequences were merged for further analyses. OBPs were also annotated in the *S. litura* genome, using a similar procedure. For OBPs and CSPs, SignalP-5.0^40^ was used to determine the presence or absence of a signal peptide. Hereafter, the abbreviations Slit, Slitu and Sfru (for *S. littoralis S. litura* and *S. frugiperda*, respectively) are used before gene names to clarify the species.

### Iterative annotation and re-annotation of gustatory receptors (GRs)

The initial annotation of gustatory receptor genes was carried out the same way as for the other genes involved in chemosensation. Subsequent steps were then added to annotate the full repertoire of GRs. At the end of the manual curation, all the newly identified amino acid GR sequences were added to the set of known GR sequences to perform a new cycle of annotation. This iterative strategy was used for *S. littoralis* as well as for *S. litura* and *S. frugiperda* and was performed until no new GR sequence was identified.

At the end of the annotation, all GR amino acid sequences were aligned for each species individually using MAFFT v7.0^41^ in order to identify and filter allelic sequences. Between alleles, only the longest sequence was retained for further analysis. Pseudogenes were identified as partial sequences containing one or multiple stop codons. Genes were considered complete when both following conditions were met: 1) a start and a stop codon were identified and 2) a sequence length >350 amino acids. *S. littoralis* gene names were attributed based on orthology relationships with *S. frugiperda* when possible. *S. frugiperda* newly identified genes compared to the previous publications were numbered starting from SfruGR232. *S. litura* newly identified gene names were numbered starting from SlituGR240.

### Annotation and enrichment analysis of transposable elements around chemosensory receptor genes in *Spodoptera* species

The annotation of transposable elements (TEs) in *S. littoralis* genome was performed using REPET (Galaxy Lite v2.5). The TEdenovo pipeline^42^ was used to identify consensus sequences representative of each type of repetitive elements. Only contigs of a length >10 kb were used as input for the pipeline. Consensus sequences were built only if at least 3 similar copies were detected in the genome. The TEannot pipeline^43^ was then used to annotate all repetitive elements in the genome using the library of TE consensus and to build a non-redundant library in which redundant consensus were eliminated (length >=98%, identity >=95%). The non-redundant library of TEs was finally used to perform the *S. littoralis* genome annotation with the TEannot pipeline.

The tool LOLA (Locus Overlap Analysis) within the R package Bioconductor^44^ was used to test for enrichment of TEs within the genomic regions containing chemosensory receptor genes (ORs and GRs) in both *S. littoralis* and *frugiperda*. To run LOLA with data from *S. littoralis*, 3 datasets were created. The first dataset, the query set, contained genomic regions of 10 kb around each chemosensory receptor gene. Since these genes were mostly organized in clusters within the genome, this dataset of the genome leaded to the creation of 114 chemosensory regions for the GRs and 63 regions for the ORs. The second dataset, the region universe, contained 1000 random regions of similar sizes selected from the genome. The last dataset, the reference dataset, contained the coordinates of TEs previously identified by the REPET analysis. The enrichment in TE content within the chemosensory regions and the control regions were then compared using LOLA using a Fisher’s Exact Test with false discovery rate correction to assess the significance of overlap in each pairwise comparison. The same method was used using *S. frugiperda* TEs, previously annotated using the same tool REPET^5^, as well as chemosensory receptor re-annotations from the present work and leaded to the creation of 191 chemosensory regions for the GRs and 88 regions for the ORs.

### Evolutionary analyses

#### Phylogenetic tree reconstructions

Chemosensory-related protein trees were constructed for OBPs, CSPs, ORs, IRs and GRs. For GRs, the phylogeny was built using GR amino acid sequences from different Lepidoptera species with various diets. In order to take into account the whole repertoire of GRs in our analysis, only species in which the GRs were annotated following whole genome sequencing were considered. The data set contained GRs from polyphagous (*S. littoralis*, *S. litura*, *S. frugiperda*), oligophagous (*H. melpomene* – 73 GRs, *Manduca sexta* – 45 GRs) and monophagous species (*B. mori* -72 GRs). The multiple sequence alignment of all GR amino acid sequences was performed with ClustalO^45^ and the phylogeny was reconstructed using PhyML 3.0^46^ (http://www.atgc-montpellier.fr/phyml/) with the automatic selection of the best substitution model by SMS^47^. The resulting phylogenetic tree was edited using FigTree v1.4.2 (https://github.com/rambaut/figtree) and Inkscape 0.92 (https://inkscape.org/fr/). Branch supports were estimated using the approximate likelihood-ratio test (aLRT)^48^ implemented at http://www.atgc-montpellier.fr/phyml/. For other gene families, sequences from various Lepidoptera species were retrieved and aligned with *S. littoralis* sequences using MAFFT^41^. The reconstruction of the phylogenetic trees was carried out the same way as for the GRs.

#### Tree reconciliation

Estimates of gains and losses of GR genes across the Noctuidae were inferred using the reconciliation methods implemented in Notung v2.6^49, 50^. The species tree was generated using TimeTree.org^51^ and the gene tree was the reconstructed phylogeny of the GRs generated by PhyML.

#### Evolutionary pressures

The codeml software of the package PAML was used to infer selective pressures^52^. Because of the high divergence between GRs across the phylogeny, selective pressures were inferred on 13 subtrees extracted from the GR phylogeny in order to minimize the ratio of synonymous substitutions. For each subtree, a codon alignment was performed using protein sequence alignments performed using MAFFT and PAL2NAL^53^ in order to convert the amino acid alignment to a codon alignment, and a phylogenetic tree was reconstructed based on this alignment. Sequences introducing large gaps in the alignment were removed in order to compute codeml on the largest alignment possible. To estimate the selective pressures acting on the evolution of the lepidopteran GR genes, the “m0 model” from codeml of the PAML package was computed on the 13 subtrees to estimate the global ω (ratio of non-synonymous substitutions dN/ratio of synonymous substitutions dS)^54^. The ω value reflects the mode of evolution, with ω>1 indicating positive selection, ω<1 indicating purifying selection and ω=1 indicating neutral evolution. To further infer positive selection, two comparisons between evolutionary models were conducted. First, the comparison between M8 and M8a models can detect positive selection acting on sites, i.e. columns of the alignment^55, 56^. This comparison was conducted only when the global ω calculated from the m0 model was > 0.3. The second comparison between branch-site model A and its neutral counterpart can detect positive selection acting on particular sites on a specific lineage^57^. Here, we tested all the terminal branches of the trees for which both the global ω was elevated and the comparison between models M8 and M8a statistically significant. Since many branches were tested for each tree, a correction for multiple testing to control for false discovery rate was applied: the q-value (Storey and al, R package version 2.22.0)^58^. In the case of a statistically significant q-value (<0.05), positively selected sites were inspected for possible artifacts due to partial sequences or misalignment.

#### Putative functional assignation

In order to assign putative functions to several candidate SlitGRs, both their phylogenetic position and theoretical 3D structure were analyzed. For the theoretical structures, the AlphaFold algorithm^59^ was used to model candidate SlitGRs as well as their *B. mori* ortholog GRs with known function: BmorGR9 and BmorGR66. Structures were then compared between orthologs using the MatchMaker tool of Chimera and the RMSD (Root Mean Square Deviation) computed using the same tool^60^.

## Results and discussion

### Genome assembly and automatic annotation of the *Spodoptera littoralis* genome

The first assembly of *S. littoralis* (v1.0), obtained with short Illumina reads, contained 123,499 scaffolds with a N50 of 18 kb and an assembly total size around 470 Mb. The second assembly (v2.0), obtained with a combination of both short Illumina and long PacBio reads, contained 28,891 scaffolds, with a N50 of 64 kb and an assembly total size around 465 Mb (Table 1). The genome size of *S. littoralis* was in good correlation with flow cytometry evaluation (470 Mb). The BUSCO analysis revealed that the second assembly contained more than 95% of complete BUSCO genes, with almost 94% being present in single-copy (Table 2). This second assembly was then used as the final assembly in all the following analyses. A total of 35,801 genes were predicted using BRAKER (OGS3.0_20171108). The BUSCO analysis indicated that almost 97% of BUSCO proteins were complete, with more than 88% being present in single-copy (Table 2). These data showed the good quality of the *S. littoralis* genome assembly, thus allowing for accurate comparison with other *Spodoptera* genomes.

**Table 1.**
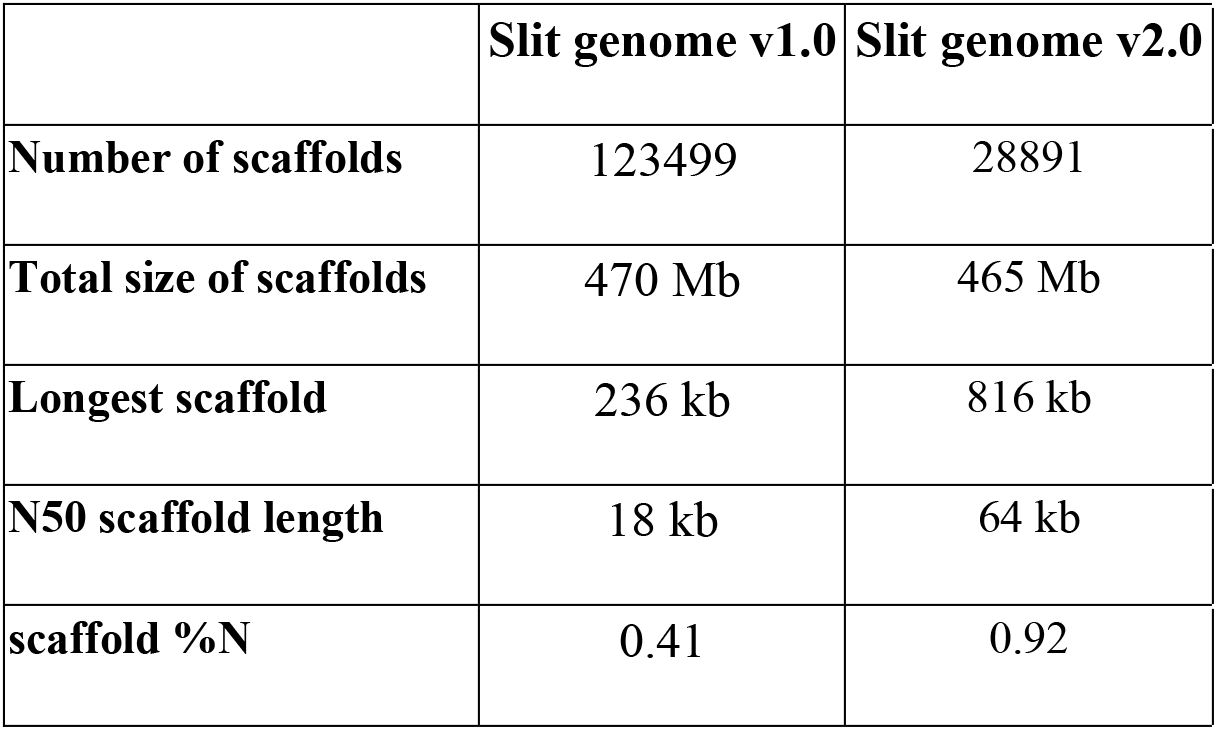
Statistics of the S. littoralis genome assemblies.

|  | Slit genome v1.0 | Slit genome v2.0 |
| --- | --- | --- |
| **Number of scaffolds** | 123499 | 28891 |
| **Total size of scaffolds** | 470 Mb | 465 Mb |
| **Longest scaffold** | 236 kb | 816 kb |
| **N50 scaffold length** | 18 kb | 64 kb |
| **scaffold %N** | 0.41 | 0.92 |

**Table 2.** BUSCO statistics on S. littoralis genome and annotation.

|  | Slit genome v2.0 | Annotation<br>BRAKER OGS3.0 |
| --- | --- | --- |
| **Complete BUSCOs (C)** | 1584 (95.5%) | 1603 (96.68%) |
| **Complete and single-copy BUSCOs (S)** | 1558 (94%) | 1467 (88.48%) |
| **Complete and duplicated BUSCOs (D)** | 26 (1.6%) | 136 (8.2%) |
| **Fragmented BUSCOs (F)** | 45 (2.7%) | 48 (2.89%) |
| **Missing BUSCOs (M)** | 29 (1.7%) | 7 (0.42%) |
| **Total BUSCO groups searched** | 1658 | 1658 |

### OBP, CSP, OR and IR chemosensory gene repertoires were of comparable size among *Spodoptera spp*

To have a full view of the *S. littoralis* chemosensory equipment, we manually curated all the major chemosensory-related gene families, including soluble carrier proteins (OBPs and CSPs), proposed to facilitate chemical transfer to chemosensory receptors^23^, and the membrane bound receptors (ORs: seven transmembrane proteins expressed in the membrane of olfactory sensory neurons, GRs: seven transmembrane proteins hosted by taste neurons, and IRs: three transmembrane proteins sensing acids and amines ^61–63^).

The genome of *S. littoralis* contained 23 CSP genes, all of them encoding full-length sequences with a signal peptide. This number of genes is similar to the 22 CSP genes annotated in *S. frugiperda*^33^ and the 23 CSP genes annotated in *S. litura*^8^. Among all these sequences, 16 CSP genes are 1:1 orthologs between the three *Spodoptera* species included in the tree while 11 CSP genes are 1:1 orthologs with BmorCSPs (from *B. mori*), showing the high level of conservation in this gene family (Supp data 2 Figure S1).

We also annotated 53 OBP genes in *S. littoralis.* Among these genes, 49 were complete and 48 possessed a signal peptide (Supp data 2). The phylogenetic tree revealed a clade enriched in *Spodoptera* OBPs (9 SlitOBPs, 9 SlituOBPs and 10 SfruOBPs) ( ). This expansion probably arose from recent tandem duplications as most of the genes of the expansion are organized in synteny in the three species (Figure S3).

We annotated 44 IR genes in the *S. littoralis* genome, 43 of which encoding a full-length sequence with various sizes containing 547 to 948 amino acids (AAs) (Supp data 2). In addition to the two conserved co-receptors IR8a and IR25a^64^, we identified 18 candidate antennal IRs putatively involved in odorant detection, 23 divergent IRs putatively involved in taste and 12 ionotropic glutamate receptors (iGluRs). The total IR number was similar to the 44 IR genes annotated in *S. litura*^65^ and the 43 IR genes annotated in *S. frugiperda*^5^. Among all these sequences, 43 IR genes are 1:1 orthologs between the three *Spodoptera* species (IR100g was missing in *S. frugiperda*). The phylogenetic tree revealed a clade containing divergent IRs and two lineage expansions were observed (IR7d and IR100), likely attributed to gene duplications^65^. The number of divergent IRs was much higher in *Spodoptera* species (*S. littoralis*: 26, *S. litura*: 26, *S. frugiperda*: 25) than in *H. melpomene* (16) and *B. mori* (6). By contrast, phylogenetic analysis (Figure S4) reveals that *S. littoralis* antennal IRs retained a single copy within each orthologous group.

We annotated 73 OR genes in the *S. littoralis* genome scattered among 61 scaffolds (Supp data 2), including the obligatory co-receptor ORco (ref Jones et al 2005 Curr Biol). The number of OR genes in the *S. littoralis* repertoire was similar to the repertoire of closely related species (69 in *S. frugiperda*, 73 in *S. litura*) and other Lepidoptera (64 in *D. plexippus*, 73 in *M. sexta*). The phylogenetic tree of ORs is presented in Figure S5.

Altogether, our annotations revealed that OBP, CSP, OR and IR repertoires were of comparable size among the *Spodoptera spp* investigated.

### A highly dynamic evolution of the GR multigene family in *Spodoptera* species

Newly obtained genomes of polyphagous noctuidae species such as *H. armigera*^66^, *S. litura*^6^*, S. frugiperda*^5^ and *Agrotis ipsilon*^67^ revealed an important expansion of gustatory receptors in these species, suggesting an adaptation mechanism to polyphagy. Here, using these known GR protein sequences and an iterative annotation process, we annotated an even larger repertoire of GRs in the *S. littoralis* genome. In view of these data, we searched for possible missing GRs in the *S. frugiperda* and *S. litura* genomes to complete their GR repertoires (Table 3). We annotated a total of 376 GR genes scattered on 110 scaffolds in the genome of *S. littoralis*, and reannotated 417 GRs on 196 scaffolds in the *S. frugiperda* genome and 293 GRs on 30 scaffolds in *S. litura* (Supp data 3). Our GR analysis not only revealed that the full repertoire of *S. littoralis* GRs is in fact much more important than previously reported, but also that the GR numbers in *S. litura* and *S. frugiperda* have been under evaluated (although the presence of some alleles may over evaluate these numbers). Among these sequences, several were indeed allelic version of previously annotated genes but several new genes were also identified (Table 3). Among these genes, the percentage of complete genes varied between species, from only 41% in *S. frugiperda* compared to 79% in *S. litura* while the percentage of complete GRs in *S. littoralis* was intermediate (73%). The percentage of allelic sequences were also highly variable, probably depending on the heterozygosity level of each considered genome. Indeed, the highest number of alleles was reached in *S. frugiperda*, a genome with a high level of heterozygosity^33^, while alleles were less frequent in the two other *Spodoptera* genomes considered. When omitting pseudogenes and alleles, the final repertoires of GRs are composed of 325 genes in *S. littoralis*, 274 GRs in *S. frugiperda* and 280 GRs in *S. litura*. As previously shown, multiple clusters of GRs were also found in the *S. littoralis* genome. The two main clusters were found on scaffolds 1414 and 878 that contained each 27 GR genes. The phylogeny reconstructed using the GR sequences from the three *Spodoptera* species as well as those from *B. mori* (BmorGRs) and *H. melpomene* (HmelGRs) showed that a few *Spodoptera* GRs clustered with candidate CO_2_, sucrose and fructose receptors, while the majority of the *Spodoptera* GRs were part of the so-called bitter receptor clades. Among the candidate bitter receptor clades, eleven clades were enriched in *Spodoptera* genes (numbered from A to K in Figure 1) and encompassed the majority of the three *Spodoptera* GR repertoires (Table 4). When belonging to the same phylogenetic clade, GRs from the same species tend to be located on the same scaffold, supporting the theory of the expansion of these genes by tandem duplications and few gene losses. For the subsequent analysis, only complete and partial genes were considered while pseudogenes were discarded. Four *S. littoralis* GRs with only one exon were identified, clustered on scaffold 67 and belonging to the same phylogenetic clade (Figure 1). Interestingly, this clade was very conserved with a 1:1 orthology relationship between the three *Spodoptera* species, the SlituGRs and SfruGRs being also monoexonic. All these monoexonic genes are orthologs with BmorGR53, a single exon gene that is highly expressed at the larval stage but not in the adult^11^. BmorGR53 is able to detect the bitter tastant and feeding deterrent coumarine^68^. It is then likely that these 4 single exon GRs play an important role in host-plant recognition in *Spodoptera* species as well.

**Figure 1.**
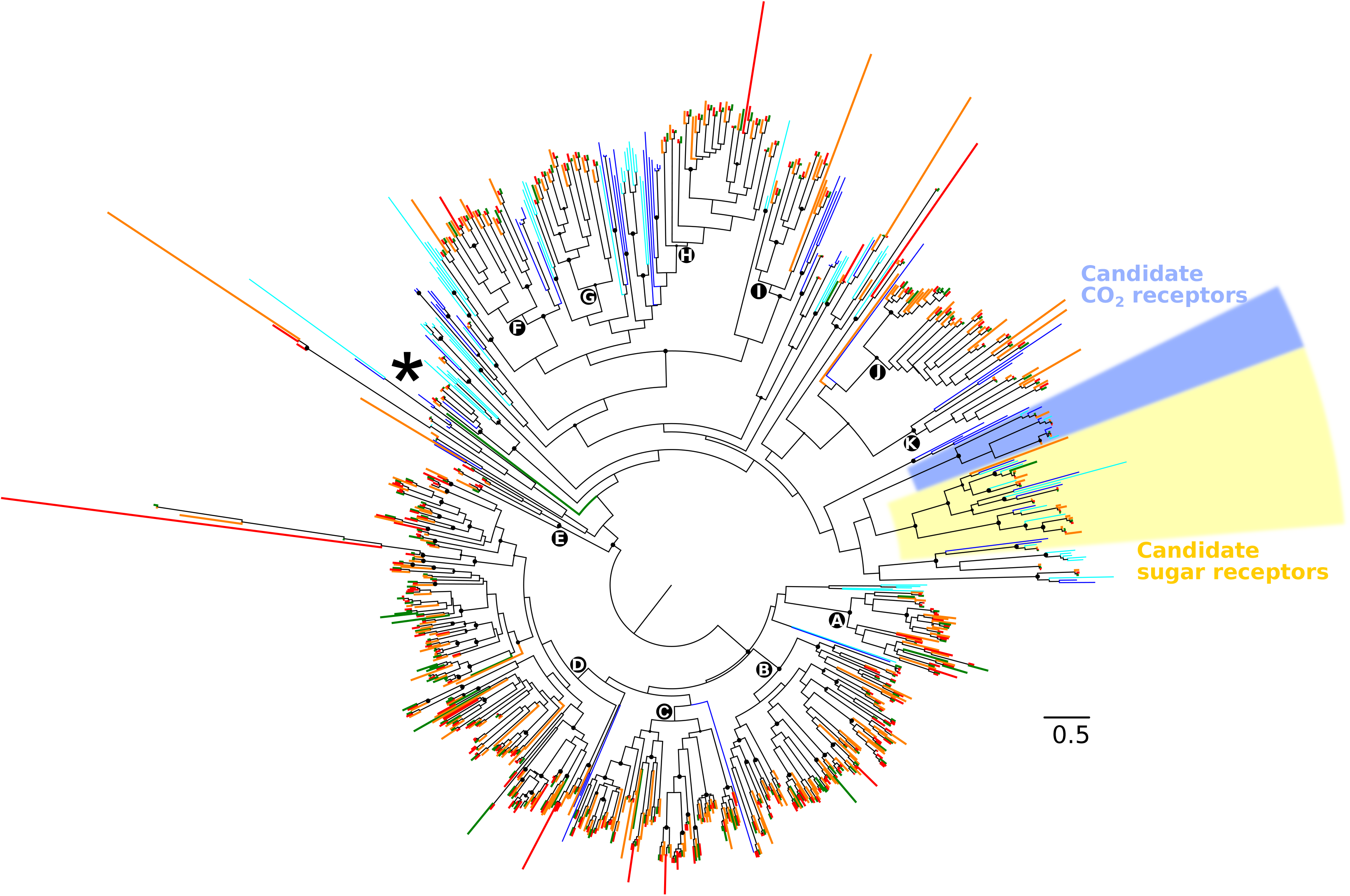
Phylogeny of lepidopteran GRs. The dataset included amino acid sequences from *S. littoralis* (Noctuoidea, red), *S. litura* (Noctuoidea, green), *S. frugiperda* (Noctuoidea, orange), *B. mori* (Bombycoidea, blue) and *H. melpomene* (Papilionoidea, cyan). Sequences were aligned using ClustalO and the phylogenetic tree was reconstructed using PHYML. CO_2_ receptor candidates as well as sugar receptor candidates are indicated in blue and yellow, respectively. All the other GRs are part of the bitter receptor clades. The star indicates the clade of single-exon GRs. Midpoint rooting was used. Circles indicate nodes strongly supported by the likelihood-ratio test (aLRT>0.9). The scale bar represents 0.5 amino acid substitutions per site.

**Table 3.** GR repertoires of *Spodoptera* species.

|  | *S. littoralis* | *S. frugiperda* | *S. litura* |
| --- | --- | --- | --- |
| **Number of GR**<br>**previously**<br>**annotated** | 38 | 231 | 237 |
| **Complete genes** | 275 (73%) | 172 (41%) | 231 (79%) |
| **Partial genes** | 50 (13%) | 106 (25%) | 49 (17%) |
| **Pseudogenes** | 19 (5%) | 22 (5%) | 7 (2%) |
| **Alleles** | 29 (8%) | 117 (28%) | 6 (2%) |
| **Total in this work** | 373 | 417 | 293 |

**Table 4.** Number of Spodoptera GRs by expansion clade. The percentages represent the proportion of Spodoptera genes to the total number of GRs annotated in the three Spodoptera species (complete + partial genes indicated in Table 3)

| **Clade** | ***S. littoralis*** | ***S. frugiperda*** | ***S. litura*** |
| --- | --- | --- | --- |
| **A** | 20 | 15 | 14 |
| **B** | 65 | 42 | 44 |
| **C** | 40 | 40 | 33 |
| **D** | 97 | 74 | 89 |
| **E** | 7 | 3 | 4 |
| **F** | 12 | 10 | 11 |
| **G** | 10 | 8 | 10 |
| **H** | 16 | 11 | 16 |
| **I** | 7 | 8 | 7 |
| **J** | 16 | 21 | 18 |
| **K** | 4 | 6 | 5 |
| **Total** | 294 (90.5%) | 238 (86.9%) | 251 (89,6%) |

The GR phylogeny served as a basis for the reconciliation of the gene- and species-tree in order to estimate gene gains and losses. The Notung analysis revealed that the ancestral repertoire of GRs of Noctuidae species contained 58 genes (Figure 2). Given the numbers of GRs annotated in *Spodoptera* species, it is not surprising that the highest gene gains occurred in the ancestor of *Spodoptera* species (296 gene gains). However, even for species with a smaller repertoire of genes such as *B. mori* (70 GRs) and *H. melpomene* (73 GRs), the turnover of genes compared to the ancestors is important (33 and 41 gene gains, 25 and 26 gene losses, respectively).

**Figure 2.**
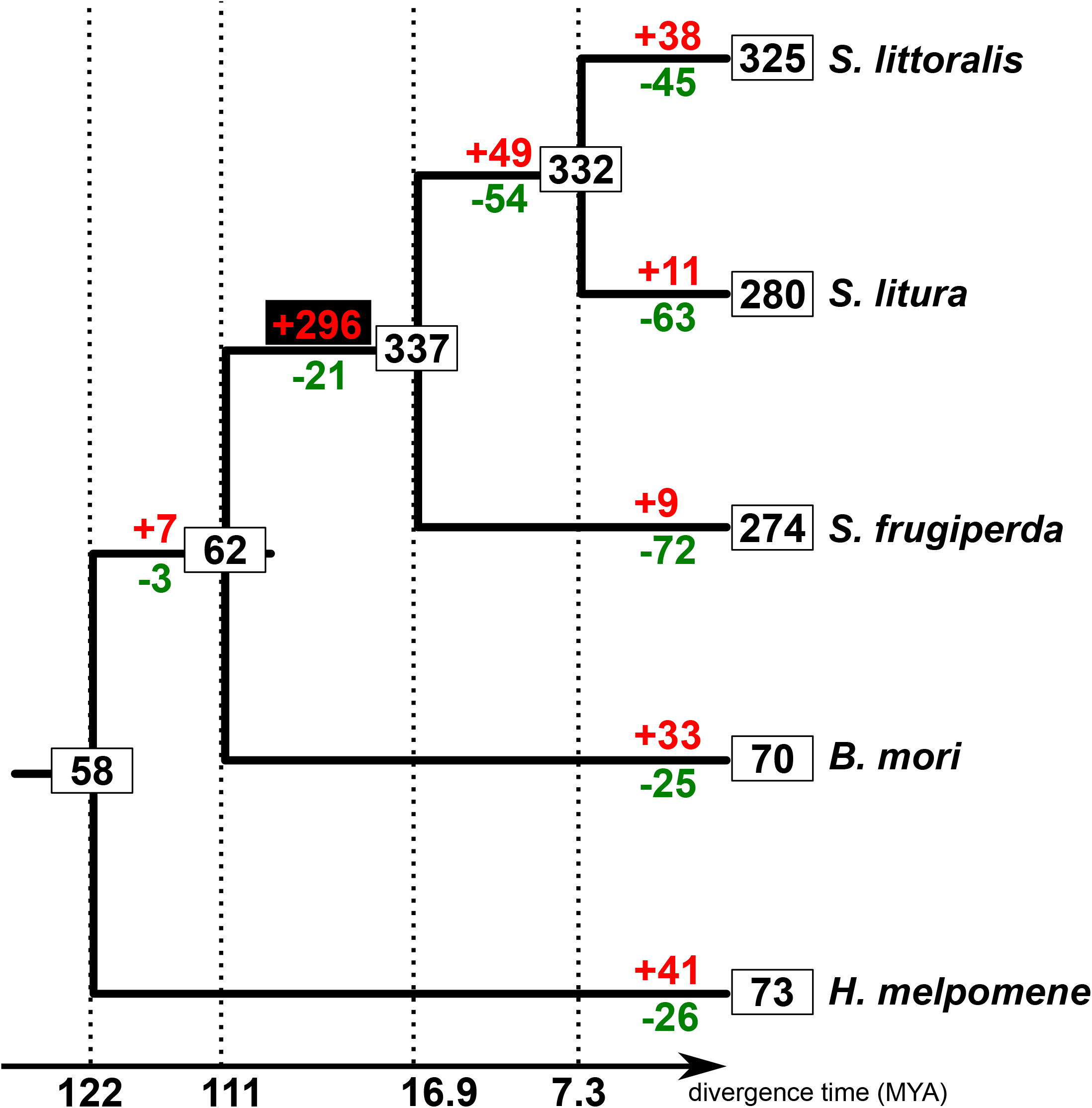
GR gain and loss estimates across lepidopterans. The gene-tree of GRs generated using PhyML was reconciled with the species-tree using Notung^50^ to estimate gene gains and losses. Numbers in boxes represent the size of GR repertoire for extant species as well as ancestors at the nodes of the species tree. Gene gains are indicated in red while gene losses are indicated in green. The expansion that occurred in the ancestor of *Spodoptera* species is indicated in red on a black background.

### Annotation of transposable elements, enrichment analysis and selection pressure

To get more insights about the mechanisms that led to the formation of massive genomic clusters of GR genes, we looked at 1) whether TEs could be involved and 2) the selective pressures acting on GR genes. TEs have been shown to be involved in countless mechanisms of evolution in insects, such as insecticide resistance, the evolution of regulatory networks, immunity, climate adaptation^16, 69–74^ and some of them have even been domesticated as genes^75^. Gene families involved in these traits have been shown to be enriched in TEs and gene family expansions have been correlated with TE content, for instance in termites^76^. Interestingly, enrichment in TEs has not been reported for insect GR gene clusters so far. While annotating GRs in the *S. littoralis* genome, we noticed the frequent co-occurrence of TEs on the same scaffolds. We thus annotated TEs in the *S. littoralis* genome and calculated their enrichment in the vicinity of GR genes. We also carried out the same enrichment analysis in *S. frugiperda* genomes, as TE annotation in this last species has been done using the same REPET pipeline as in our study. The *de novo* constructed library contained 1089 consensus sequences of TEs and was used to annotate the *S. littoralis* genome. The repeat coverage for the *S. littoralis* genome was 30.22%, representing 140 Mb, which is similar to that of *S. frugiperda* (29.10%), *S. litura* (31.8%) and *S. exigua* (33.12%)^5, 6, 77^. The relative contribution of the different classes of repetitive elements revealed that Class I elements were more represented than Class II elements (66.96% vs 20.83%), a classical feature of insect genomes^75^ (Figure 3, Table 5). However, the repartition and proportions between the different classes differed between these species. The Class I SINE was the most represented in *S. frugiperda* (12.52%)^33^ and one of them was found to be enriched in the vicinity of the GRs in both *S. littorali*s and *S. frugiperda* while the class I LINE elements were the most represented in both *S. litura* and *S. exigua*, although with a lower proportion of all repeated elements (27.73% and 14.81%, respectively). Remarkably, the proportion of LINE elements identified in the *S. littoralis* genome was the highest reported so far in arthropods^78^, accounting for 52.18% of all repetitive elements. In two subspecies of the Asian gypsy moth *Lymantria dispar*, the accumulation of this particular class of transposable elements was found to be responsible for their large genome size^79^, a phenomenon also observed in other insect species^75^. The accumulation of the same elements in the *S. littoralis* genome could explain its larger size compared to its *Spodoptera* counterparts (470Mb vs ∼400Mb for *S. frugiperda*, 438Mb for *S. litura*, 408-448Mb for *S. exigua*). The second most represented was DNA transposons, Class II TIR elements, representing 11.04% of all TEs (Figure 3, Table 5).

**Figure 3.**
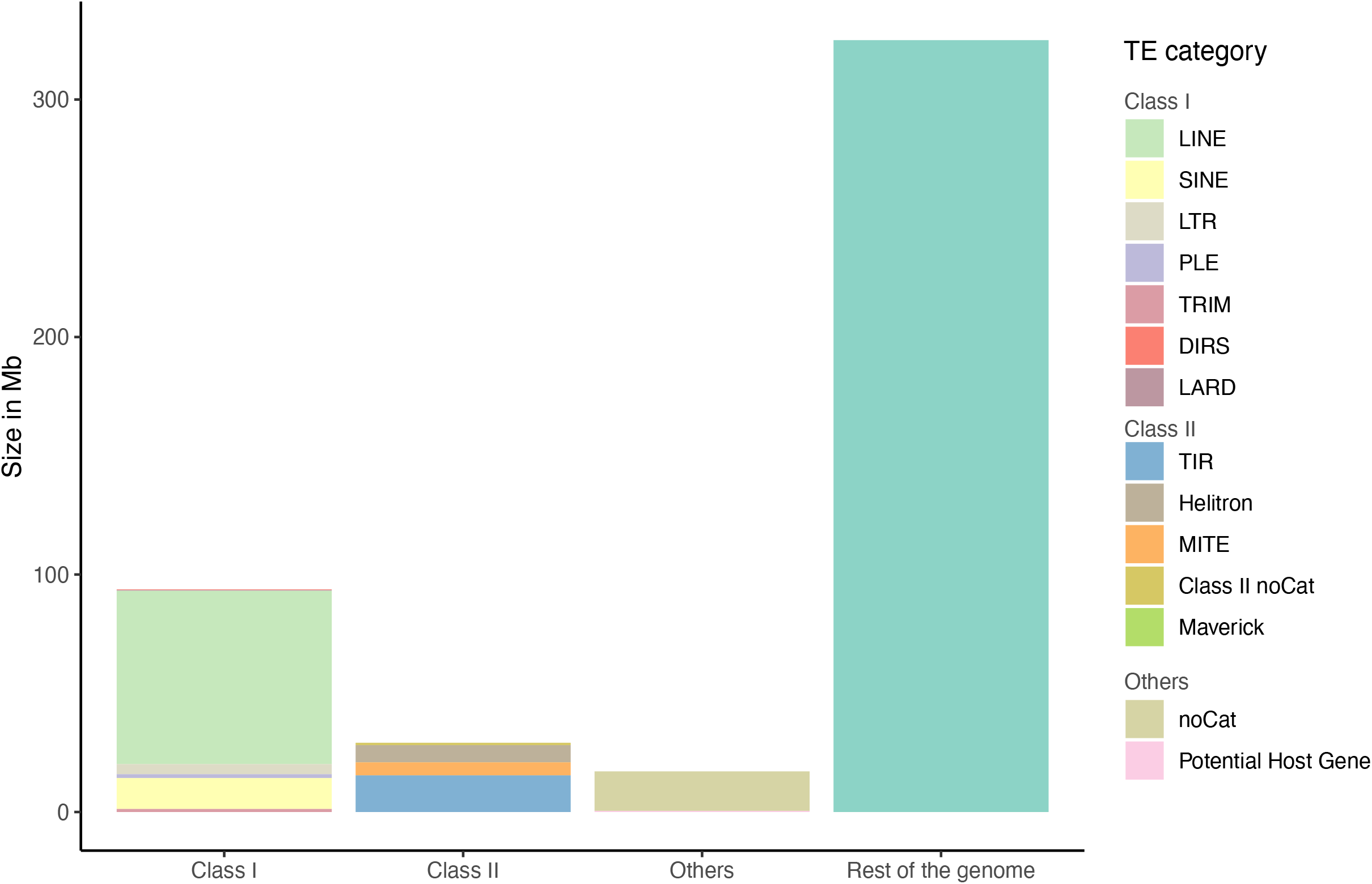
Repartition and size of repeat content in *S. littoralis* genome. Repetitive elements account for 30.22% of *S.littoralis* genome. Class I elements are more abundant than class II. The class I LINE elements represent more than half of all repetitive elements.

**Table 5.** Repartition of repetitive elements in *S. littoralis* genome based on the classification established by Wicker et al.^97^. noCat means repetitive elements that could not be classified into the existing categories.

|  | TE category | % of coverage of all repetitive elements |
| --- | --- | --- |
| Class I Retrotransposons | DIRS | 0.20% |
|  | LARD | 0.19% |
|  | **LINE** | **52.18%** |
|  | LTR | 3.02% |
|  | PLE | 1.12% |
|  | SINE | 9.33% |
|  | TRIM | 0.92% |
| Class II DNA transposons | Helitron | 5.17% |
|  | MITE | 3.90% |
|  | Maverick | 0.01% |
|  | **TIR** | **11.04%** |
|  | Class II noCat | 0.71% |
| Others | noCat | 11.87% |
|  | Potential Host Gene | 0.35% |

The enrichment of TEs in the vicinity of GR gene clusters was tested in both *Spodoptera littoralis* and *frugiperda*, and we found that a particular category of class I retrotransposons, a SINE transposon, was significantly enriched (q-value < 0.005) in the vicinity of the GRs in both species (q-value = 0.0043 and 0.0078, respectively), suggesting a transposon-mediated mechanism for the formation of GR clusters (Supp data 4). SINE elements are typically small (80-500 bp) and originate from accidental retrotransposition of various polymerase III transcripts. These elements are non-autonomous, therefore their involvement in the dynamic of the GR multigene family may be related to their potential to induce genome rearrangements via unequal crossing over, hence potential drivers of duplication, as previously shown in other insect species ^80–83^. Given their prevalence in the *S. littoralis* genome, the potential role of these TEs in the GR family dynamic is probably just one of their numerous functions. The same enrichment analysis performed for the OR loci showed no significant enrichment in *S. littoralis* but did show enrichment in an uncharacterized class of transposons in the vicinity of SfruORs (Supp data 4).

Several studies have shown the importance of positive selection in the evolution of multigene families, especially in chemosensory genes such as ORs and GRs^84, 85^. Positively selected chemoreceptors may be linked to adaptation in *Drosophila* species^86, 87^. In the pea aphid, signatures of selection have been identified in chemosensory genes, including GRs and ORs, which may be implicated in the divergence of pea aphid host races ^88–90^. We thus analyzed selective pressures focusing on 13 clades of interest in the *Spodoptera* GR phylogeny: the potential clade of CO_2_ receptors, the potential clade of sugar receptors and the 11 expended lineages within the so-called bitter receptor clades. For all 13 clades, we observed low global ω values ranging from 0.01 to 0.42, with the highest observed for candidate bitter receptor clades. The comparison between models M8 and M8a was statistically significant for clades C, F and J, indicating a signal of positive selection. Branch-site models on terminal branches of the associated trees were then tested on these clades. For clade J, no GR was revealed as evolving under positive selection. However, for clades C and F, 2 and 5 GRs were identified as positively selected, respectively (Table 6). Within these GR sequences, very few positively selected sites were identified for each gene (between 0 and 3, Supp data 5). This finding is coherent with previous studies showing the same pattern of evolutionary rates^91, 92^, especially in *S. frugiperda*^5^ (3 GRs under positive selection when comparing two host strains). Taken together, all positive selection analyses indicate that positive selection within the GR gene family is cryptic and may not play an important role in shaping the evolution of *Spodoptera* GRs. Anyhow, the few positively selected GRs may be interesting candidates for further functional studies.

**Table 6.** Selective pressure analysis. NS: non significant, /: not calculated.

| Clade | # of sequences | $\omega$ (M0) ($d_N/d_S$) | p-value (M8 vs M8a) | branch-site |
| --- | --- | --- | --- | --- |
| **A** | 45 | 0.34109 | NS | / |
| **B** | 133 | 0.34146 | NS | / |
| **C** | 86 | 0.34386 | 0.044804* | Slit_GR217, Slitu_GR155 |
| **D** | 218 | 0.29174 | / |  |
| **E** | 14 | 0.18639 | / | / |
| **F** | 33 | 0.41616 | 0.000504** | Sfru_GR44, Sfru_GR49,<br>Slit_GR44 |
| **G** | 28 | 0.36571 | NS | / |
| **H** | 38 | 0.31933 | NS | / |
| **I** | 16 | 0.22375 | / | / |
| **J** | 38 | 0.42257 | 0.005319** | NS |
| **K** | 11 | 0.17393 | / | / |
| **Sugar** | 27 | 0.05662 | / | / |
| **CO2** | 11 | 0.01074 | / | / |

### Putative functional assignation of candidate SlitGRs

The complexity of the evolution of the bitter GRs is reflected by their complex functioning. Indeed, in contrast with the relatively simple OR/Orco association that is the basis for olfaction, the molecular basis for gustation is marked by several characteristics that were recently identified in *D. melanogaster*. First, some GRs have to be co-expressed within the same neuron in order to be able to respond to a stimulus. Second, it seems that GR-GR inhibition can modulate neuron responses ^93^. The challenge in the next few years will be to characterize both the response spectra and precise expression patterns of GRs of interest. However, those GRs of interest need to be selected. The present work provides us with some valuable candidates such as the single exon GRs for which the function is known in *B. mori*. Also, it seems that individual GRs can play an important role in the ecology of a species. Among examples are BmorGR9, which binds D-fructose without the need of any other GR^94, 95^, and BmorGR66, whose silencing confers to the monophagous *B. mori* larva the ability to feed from different food sources^96^. We identified their *S. littoralis* orthologs as SlitGR9 and SlitGR15, respectively. We predicted their 3D structures using AlphaFold and compared them to the AlphaFold predicted structures of *B. mori* orthologues. The RMSD computed between the whole 3D structures of BmorGR9 and SlitGR9 was 10.855Å (Figure 4A), but when the N-terminal end, as well as the loop between the transmembrane domains 4 and 5, were removed (regions that were disordered and difficult to predict), the RMSD improved to 1.170Å (Figure 4B). Both structures were strikingly similar on their extracellular side, suggesting that SlitGR9 is likely a D-fructose receptor in *S. littoralis*. The ligand of BmorGR66 is not known, however, this receptor is responsible for the feeding difference of *B. mori* for mulberry leaves^96^. Its ortholog SlitGR15 is a key candidate for functional studies to test if this GR has an impact on the feeding preference in *S. littoralis* as well. When comparing both full structures, the RMSD was 6.518Å (Figure 4C) while it decreased to 3.599Å when the N-terminal of both structures were removed from the analysis (Figure 4D). Interestingly, differences were visible between both structures in the extracellular domains of the proteins, suggesting that the binding pockets may differ as well.

**Figure 4.**
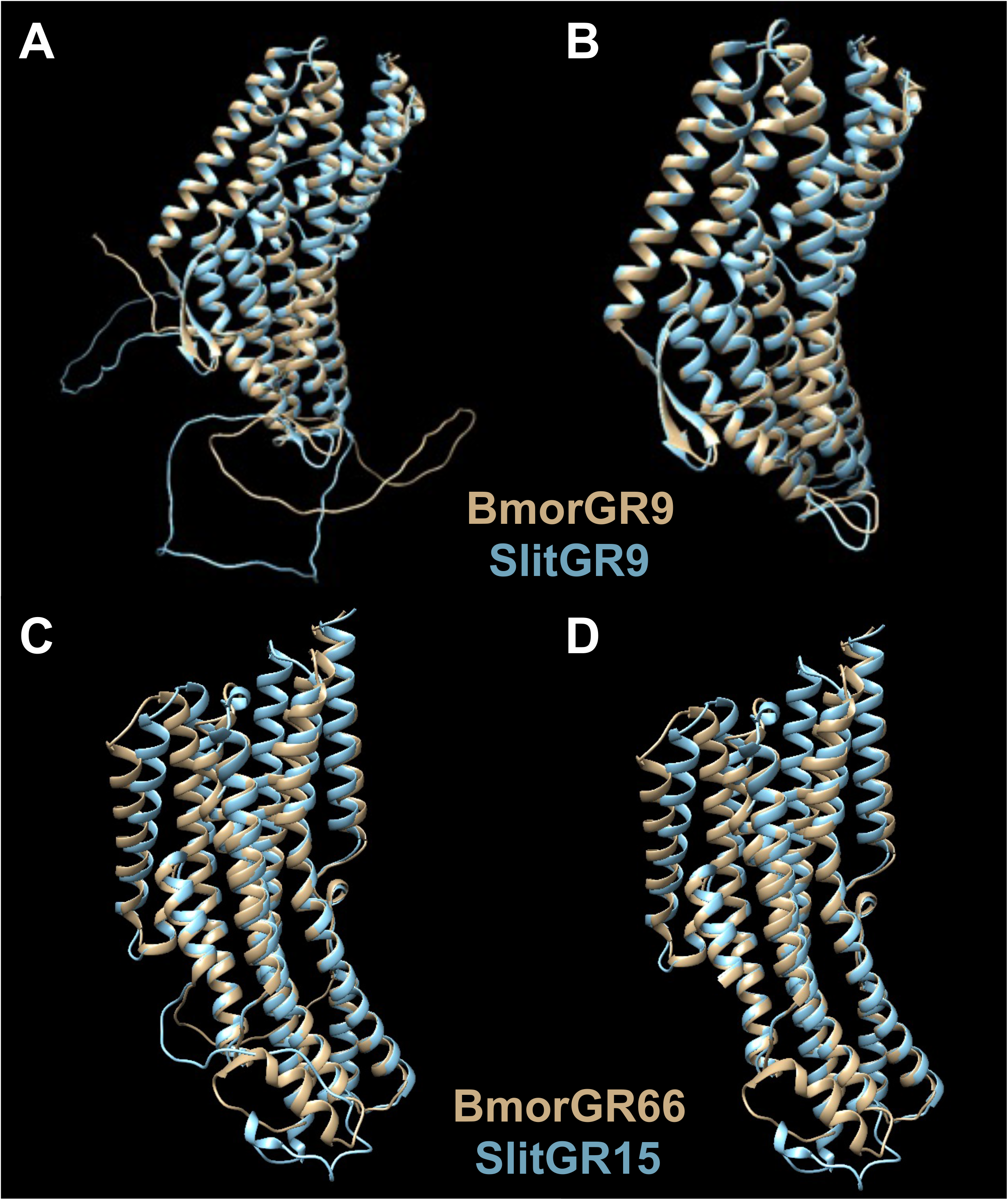
3D structures of BmorGR9 and BmorGR66 and their respective orthologs, SlitGR9 and SlitGR15. 3D structures were predicted using AlphaFold2^59^. **A** and **C**. Alignment of full 3D structures. In **B** and **D**, the disordered regions such as the large extracellular loops (**B**) and the N-terminal ends (**B** and **D**) were removed for the comparison of the orthologs’ structures.

Apart from these particular GRs, the neuronal coding of taste via more than 200 genes in species like *Spodoptera* moths is not known. But are all these GRs at play in effective taste sense? In fact, comparison of GR gene repertoires with transcript repertoires showed that a small proportion of the gene repertoire is actually expressed in the canonical gustatory tissues of *Spodoptera spp*., as can be seen in *S. littoralis* and *S. litura*^8, 20, 21^. In addition, GR expression levels - especially that of bitter receptors - are rather low. Whether the genome acts as a “reservoir” for a multitude of GR genes to be selectively expressed in accordance with the evolution of food preference remains to be investigated. In that view, the identification of regulatory genomic regions and transcription factors in the vicinity of GR regions that may be at play in GR expression choice would help understanding if and how GRs evolved according to polyphagy.

## Supplementary figures

**Figure S1.**
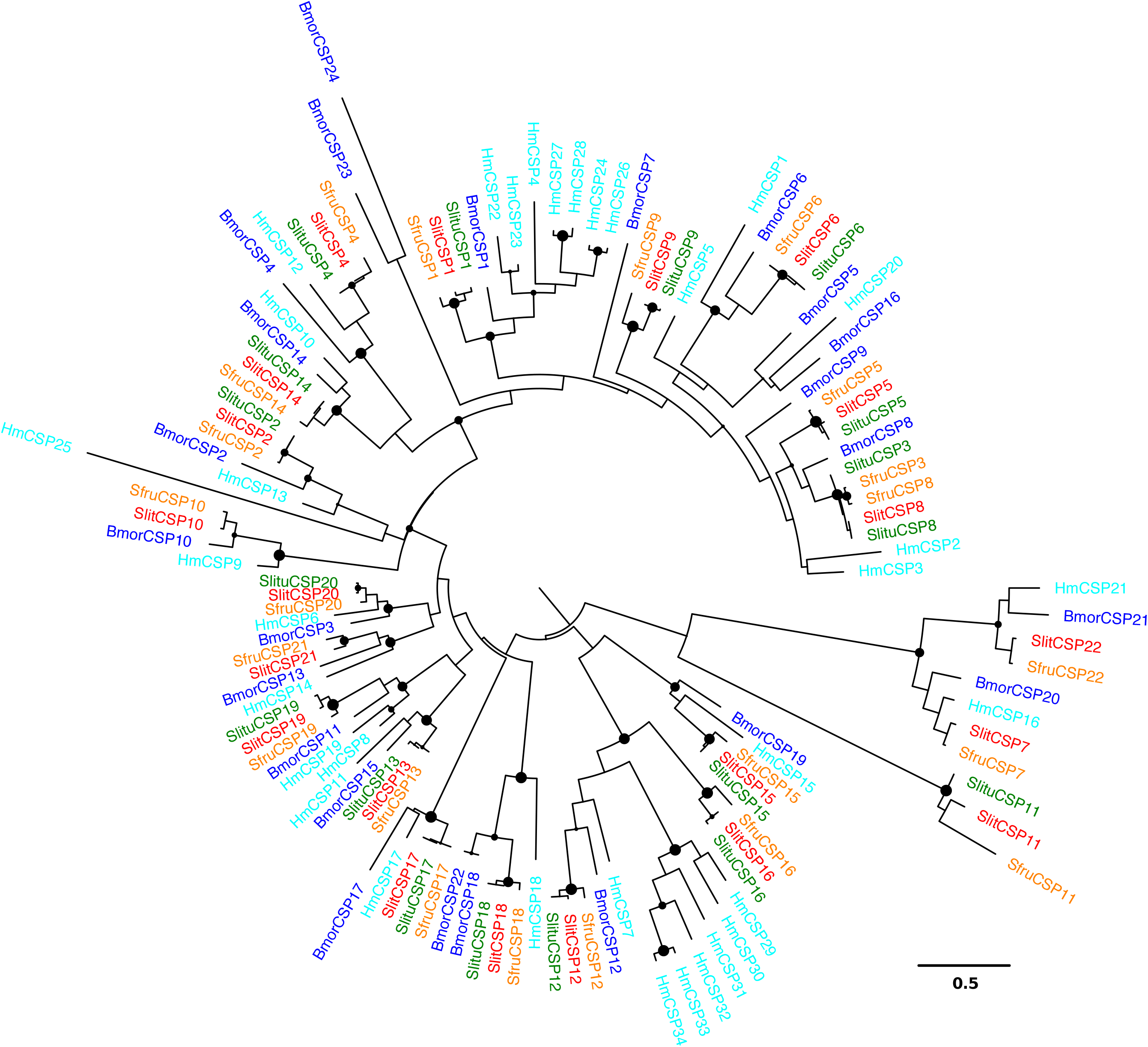
Phylogeny of lepidopteran CSPs. The dataset included amino acid sequences from *S. littoralis* (Noctuoidea, red), *S. litura* (Noctuoidea, green), *S. frugiperda* (Noctuoidea, orange), *B. mori* (Bombycoidea, blue) and *H. melpomene* (Papilionoidea, cyan). Sequences were aligned using MAFFT and the phylogenetic tree was reconstructed using PhyML. Midpoint rooting was used. Circles indicate nodes strongly supported by the likelihood-ratio test (aLRT>0.9). The scale bar represents 0.5 amino acid substitutions per site.

**Figure S2.**
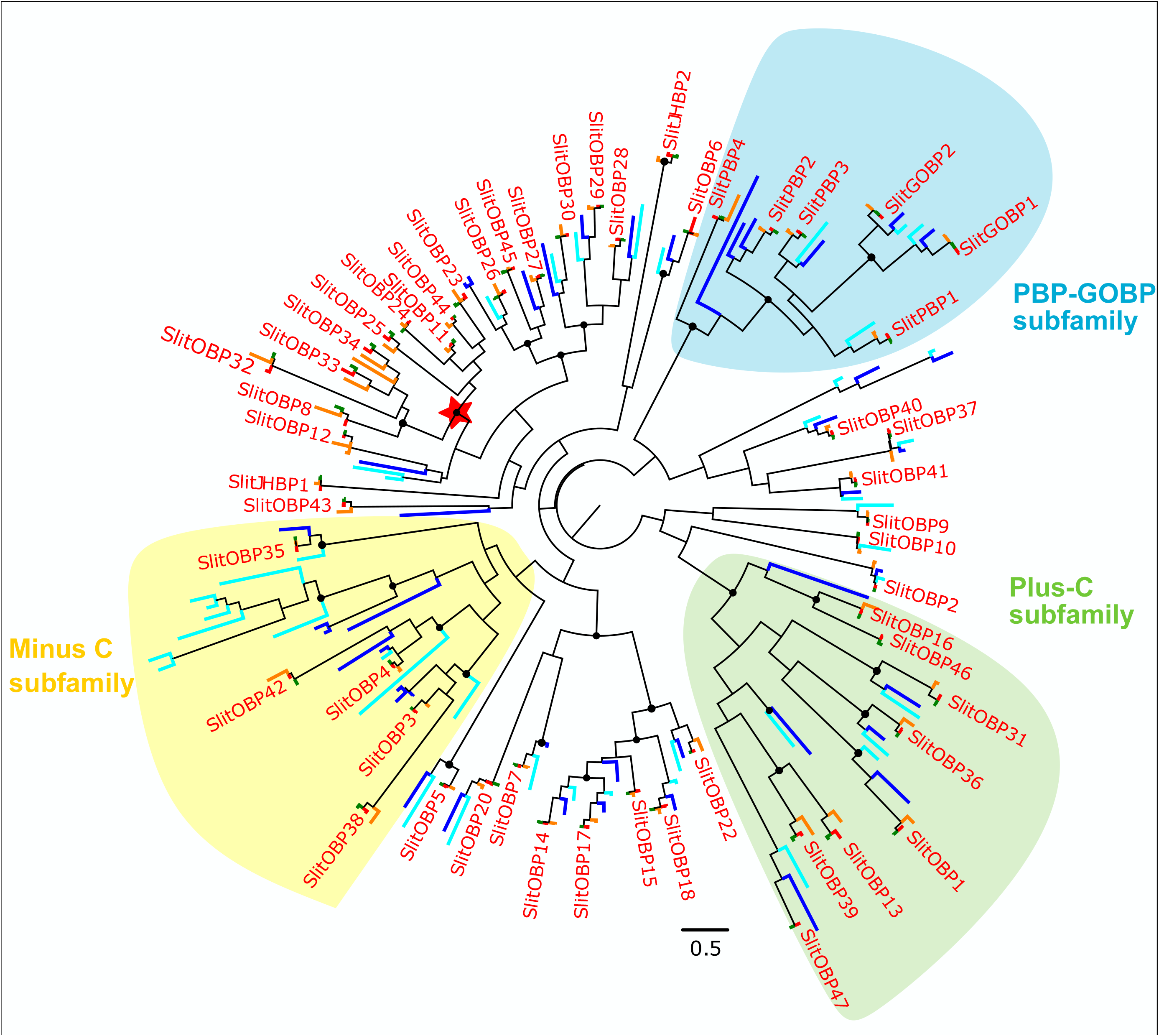
Phylogeny of lepidopteran OBPs. The dataset included 53 amino acid sequences from *S. littoralis* (Noctuoidea, red), 53 sequences from *S. litura* (Noctuoidea, green), 53 sequences from *S. frugiperda* (Noctuoidea, orange), 44 sequences from *B. mori* (Bombycoidea, blue) and 43 sequences from *H. melpomene* (Papilionoidea, cyan). Sequences were aligned using MAFFT and the phylogenetic tree was reconstructed using PhyML. Subfamilies are indicated with different colors (yellow: Minus C subfamily, green: Plus-C subfamily, blue: PBP-GOBP subfamily). Midpoint rooting was used. Circles indicate nodes strongly supported by the likelihood-ratio test (aLRT>0.9). The red star indicates expansion in *Spodoptera*. The scale bar represents 0.5 amino acid substitutions per site.

**Figure S3.**
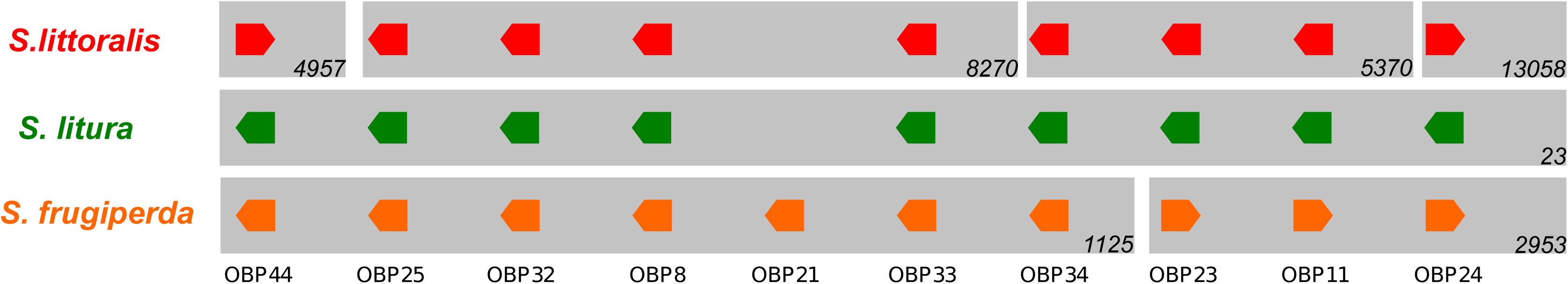
Genomic organization of the *Spodoptera* OBP genes. Scaffolds/chromosomes are represented in gray, with their numbers in italic. Gene names are indicated and their orientations are represented by the arrows.

**Figure S4.**
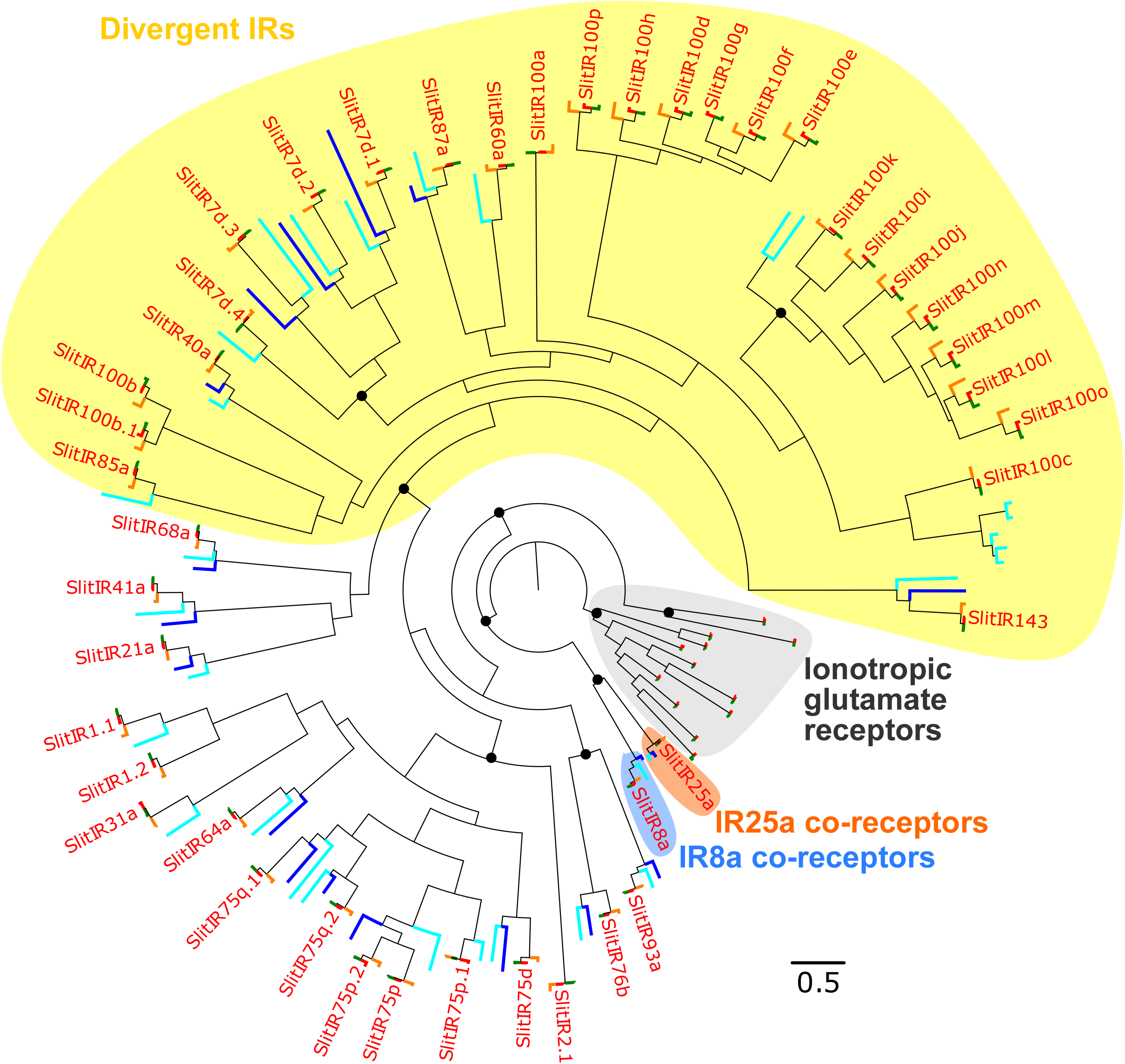
Phylogeny of lepidopteran IRs. The dataset included amino acid sequences from *S. littoralis* (Noctuoidea, red), *S. litura* (Noctuoidea, green), *S. frugiperda* (Noctuoidea, orange), *B. mori* (Bombycoidea, blue) and *H. melpomene* (Papilionoidea, cyan). Sequences were aligned using MAFFT and the phylogenetic tree was reconstructed using PHYML. Colors indicate different categories of IRs (yellow: divergent IRs, grey: ionotropic glutamate receptors, orange: IR25a coreceptorsThe tree was rooted using the iGluR clade. Circles indicate basal nodes strongly supported by the likelihood-ratio test (aLRT>0.9). The scale bar represents 0.5 amino acid substitutions per site.

**Figure S5.**
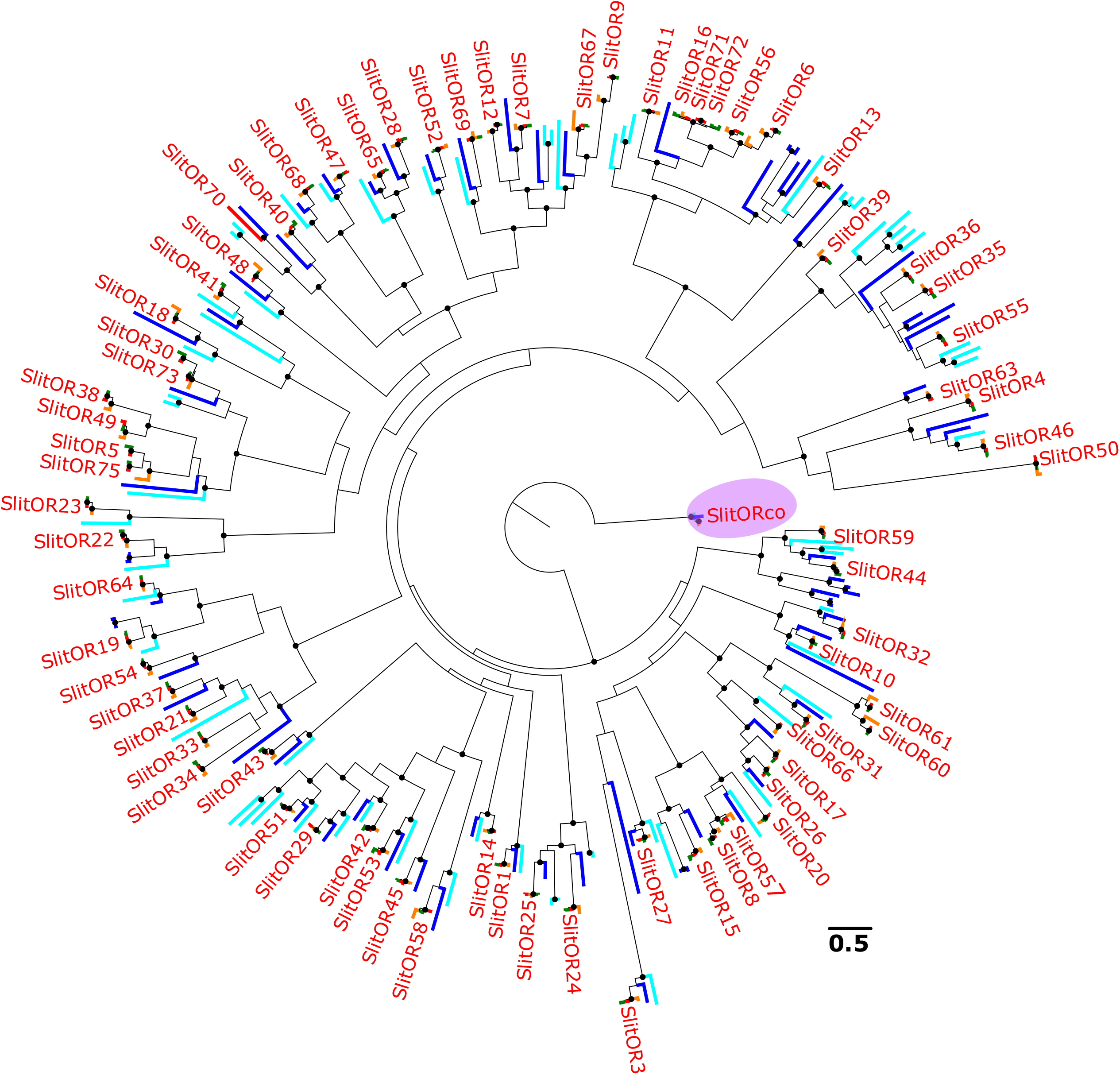
Phylogeny of lepidopteran ORs. The dataset included amino acid sequences from *S. littoralis* (Noctuoidea, red), *S. litura* (Noctuoidea, green), *S. frugiperda* (Noctuoidea, orange), *B. mori* (Bombycoidea, blue) and *H. melpomene* (Papilionoidea, cyan). Sequences were aligned using MAFFT and the phylogenetic tree was reconstructed using PHYML. The tree was rooted using the Orco clade (purple). Circles indicate basal nodes strongly supported by the likelihood-ratio test (aLRT>0.9). The scale bar represents 0.5 amino acid substitutions per site.

## Data Availability Statement

The assembled genome of *Spodoptera littoralis* as well as the genomic data of *Spodoptera litura* (v1.0)^8^ and *Spodoptera frugiperda* (Corn variant, v3.1)^33^ are all publicly available on the BIPAA platform (https://bipaa.genouest.org) and on NCBI (XXXX).

## Acknowledgments

The *S. littoralis* genome has bseen sequenced in the framework of the i5K initiative (http://i5k.github.io/) and the InsectGenomes projectprojet at INRAE leaded by Denis Tagu (IGEPP).

