## Supplementary material for "*Spodoptera littoralis* genome mining brings insights on the dynamic of expansion of gustatory receptors in polyphagous noctuidae": Supp_data5

BEB prob (ω>1) > 95%

BEB prob (ω>1) > 99%

CladeC

>Slitu_GR155

MFWHLTSTKKRPVKIYKHKILPYEEVLLNNVIEKDLQSILRPLNFMHHCFVCAKYTIRDNFITSNSLKYKLFGLICALFYRFFLLYDFGRNIYLYWNKFLTTLLIRLGYQNVTLAIGYFLIFFSNFLYCNDNVVLVVNIQNLIRTFKLERKHLNSFIIFNWFWVIFINFIFIFENYRNIITIDFRNFIFINVFSSIPSILYDINIVYAIFFVNLLKKTLKIMTKDLVRSSVRVSNSRSHWTEFYNSFANILETYNLFQKTFRLLIAFYSFFTISNSLLNVSIFITLGHDFNVIHITDILITFCSFVGRHILLLTLLCVQSEKLYAAFDESSYNSNLLCSLSESQRIVCRNIQRLYKASFKRFTVYGMFVIDVNFLFHLVAVISMYTIAQLQLILPPDE

>Slit_GR217 (no sites)

CladeF

>Sfru_GR44

MDVILKDVLNYKSNVRKFINISISILRFAVGNYKKFNDSKYICFLAKLYCITVACCIIFRQIHVHGIFSLTKVIGRNPTRSRTRMKMLSNERYMERFSKGLSTCDAIMGFKDKSIMTEILFGISVSILIAKGLISLFWCYNISFELSTCVMMFSTEFNYLLHCVCQMTLYNRMGFIKKRLQSNLIHINIVGKDEIGRNVRVVRKCLGYYHNLLDNVQQLDTAMQVLVMALVISMMHVLAPSFITEMVNNNIDDIKSTLVTQMVRCSDKSLREELDTALQYIHRRPYKFVICGAVAVDGKLPFSIIGVCITYVILIKQFTHFVDVTS

>Sfru_GR49

MYHFKHFIQRHTPSNWFLKLLVILRLLLTNYTDISESRAKCYLLKFYCISIWFCFSLFYFYDVFARNAFGIYSLCVVEYTLCGIANFIGGDDFFFKFIRVIEINDRIIGFKKMSFFTKYLCVITVSNVTIRLFISITHAIVDPRPQYLFATVTFALLSTDINHIFNILIFSIIQNRMKQLQLFFKSIYIPVNISGRNEVEINIKSIRKGLLYYNNLLDNLRSISKSLQVLLFINWIIHCVKTLFLGFIIASLISRVHHKQLLGMMMEVMQSLIMISSPAIIASITANHVNDIKRMLSSKLLQTSDESLVFELETTLQYMTLRPFQXVVTLNIYLPFVVIGLCITYVVVGLQLSQIKI

>Slit_GR44

MDGFLKNLLSYKSNVRKMYIITITVLRFVVGNYQKFNDSKRLCFFSKLYCITVACCLIYSHLNIHGLCFSPNLFAIEYSIYFVYYFITGQEYLMKFSKGLSTCDAIMNFKDMSILTEILFGISVSVLTAKYLTLYLYYNLSFDLSTCTVMFSTVFNYLLNYVLQMTLYNRMRIIKKCLQSNVIHINIVGKDQIGRNVRVVRKCLRYYHNLLDNIQELDITMQVLLSASLLCNIPQWIGGFSLSTDLWFNNVSGSVDLPMSMIHVMAPAIITELINNKIDEIKSTLVTQMIRCSDKSLREELETALQYIRRRPYKFVICGAVTVDGRLPISILSICITYVILTKQFIHFVDITI

>Slit_GR49 (no sites)
